# D-Methionine Improves Spatial Navigation and Attenuates Oxidative Stress and Amyloid Pathology in a Sex-Specific Manner

**DOI:** 10.64898/2026.01.27.702104

**Authors:** Mackenzie R. Peck, Jenelle E. Chapman, Tiarra Hill, Kathleen Quinn, Erol D. Ikiz, Angel Lopez, Erin R. Hascup, Chilman Bae, Kevin N. Hascup

## Abstract

**Background:** Oxidative stress and maladaptive neuroimmune activation contribute to cognitive decline in Alzheimer’s disease (AD) and represent therapeutic targets beyond amyloid-centered approaches.

**Objective:** To determine whether oral D-methionine (D-Met), a redox-active amino acid, reduces amyloid pathology and lipid peroxidation and confers disease-modifying benefits in AD mouse models.

**Methods:** Male and female APP/PS1 and APP^NL-F^ mice with advanced AD pathology received oral D-Met or vehicle. Behavioral assessments included locomotor activity and hippocampal-dependent spatial learning and memory. Amyloid burden, lipid peroxidation, peripheral metabolic and inflammatory markers, and hippocampal microglial phenotypes were evaluated using biochemical and histological analyses.

**Results:** D-Met did not alter locomotor or exploratory behavior but improved spatial memory recall in both sexes of APP/PS1 mice and in female APP^NL-F^ mice. APP^NL-F^ males exhibited improved learning during Morris water maze (MWM) acquisition. Amyloid pathology was modestly and region-specifically reduced, including decreased hippocampal plaque size in male APP^NL-F^ mice, reduced cortical plaque size in female APP/PS1 mice, and lower soluble amyloid-β (Aβ)_42_ in male APP/PS1 mice. Lipid peroxidation, assessed by malondialdehyde, was reduced only in female APP^NL-F^ mice. D-Met induced pronounced sex-dependent peripheral effects, increasing adiposity and pro-inflammatory adipose signaling in males, while reducing perigonadal white adipose tissue (pgWAT) IL-6 expression in female APP^NL-F^ mice. In the hippocampus, D-Met remodeled microglial signatures, with female APP^NL-F^ mice showing reduced Iba1 and disease-associated microglial (DAM) markers and increased Axl expression.

**Conclusion:** Short-term D-Met acts as a metabolic and redox modulator with modest amyloid-lowering effects mediated by improved microglial function. Therapeutic efficacy is strongly sex- and model-dependent, with the greatest benefit observed in female APP^NL-F^ mice.

## Introduction

AD is an age-related neurodegenerative disorder characterized by progressive accumulation of Aβ plaques and neurofibrillary tangles. These pathological hallmarks cause a cascade of neurobiological changes resulting in progressive deficits in new learning and memory, neurodegeneration, and eventual death. Until recently, the limited medications available to treat AD were primarily symptomatic. Current FDA-approved monoclonal antibody treatments aid in plaque removal, and offer modest disease-modifying effects, yet are associated with risks such as edema or microhemorrhages. This suggests additional research is needed to identify safer alternatives that modulate endogenous clearance mechanisms that have the potential to provide disease-modifying benefits.

Microglia are specialized macrophages of the CNS that can surround plaques and facilitate their clearance through phagocytic mechanisms ^1^. Oxidative stress and lipid peroxidation are central features of AD that compromise microglial capacity to clear Aβ aggregates thereby exacerbating plaque accumulation and neurodegeneration. This phagocytic impairment occurs through several mechanisms including reactive oxygen species (ROS) production that oxidatively modifies cell surface receptors and signaling pathways essential for engulfment and motility ^2^. Oxidative environments also shift microglial phenotypes toward pro-inflammatory states that produce cytokines leading to neuroinflammation. The subsequent oxidative and inflammatory stress leads to accumulation of lipid droplets within microglia and inversely correlates with their Aβ phagocytic capacity ^3^. Altogether, this creates a feed-forward cycle of hindered phagocytic clearance and chronic inflammation leading to plaque accumulation.

Plaques are more rigid than the surrounding parenchyma, and microglial cells detect and migrate towards this stiffness through activation of plasma membrane-bound Piezo1 channels ^4^. Piezo1 receptors are mechanosensitive nonspecific cation channels that transduce mechanical stimuli into cellular signaling mechanisms. Recent studies have shown that Piezo1 knockout impairs phagocytic clearance of Aβ plaques while chemical activation by Yoda1 reduced pathological burden in 5xFAD mice ^5^. Although agonists such as Yoda1 exhibit strong specificity for Piezo1 receptors, their activation initiates a signaling cascade that upregulates P2X3 receptors, thereby sensitizing sensory nerve fibers and resulting in hyperalgesia or allodynia. ^6,7^. These side effects limit the translational applications of Piezo1 agonists and suggest receptor modulators are needed for therapeutic AD interventions.

D-Met exerts otoprotective effects from cisplatin ^8^ and noise-induced hearing loss ^9^ through free radical scavenging and prevention of lipid peroxidation. Recently, D-Met has been shown to modulate Piezo1 channel kinetics by delaying receptor inactivation (unpublished observations). Considering the influence of Aβ_42_-mediated oxidative stress ^10^ and the neuroprotective function of microglial cells, D-Met administration may provide a dual therapeutic approach for treating AD. We hypothesized that D-Met would reduce Aβ plaques and lipid peroxidation thereby providing disease-modifying therapeutic benefits in AD mouse models. We tested this hypothesis in the transgenic APP/PS1 and knock-in APP^NL-F^ AD models. Transgenic APP/PS1 mice overexpress amyloid precursor protein (APP) with preferential cleavage for the Aβ_42_ isoforms leading to prevalent plaque deposition throughout the neocortex and spatial navigation deficits by 12 months of age ^11,12^. APP^NL-F^ mice overproduce Aβ_42_ without overexpression of the APP leading to slower pathological progression and spatial navigation deficits by 18 months of age ^13^. Accordingly, to evaluate the potential disease-modifying effects, we initiated D-Met treatment at ages corresponding with plaque burden and memory recall deficits of each model. We found that 1 month of oral D-Met treatment modestly improved spatial navigation performance through sexually dimorphic mechanisms.

## Methods

### Animals

Male and female AβPPswe/PSEN1ΔE9 (APP/PS1) and APP^NL-F^ mice expressing the Swedish (NL) and Iberian (F) mutations were used for all assays. Mice were bred and maintained in our animal colony and originated from founder C57BL/6J (RRID:IMSR_JAX:000664) and APP/PS1 (RRID:MMRRC_034832-JAX) mice from Jackson Laboratory (Bar Harbor, ME). Our colony of APP^NL-F^ mice on a C57BL/6J background (RRID: IMSR_RBRC06343) were obtained from Riken (Japan). Protocols for animal use were approved by the *Institutional Animal Care and Use Committee* at Southern Illinois University School of Medicine (Protocol number: 2025-055), which is accredited by the Association for Assessment and Accreditation of Laboratory Animal Care. All studies were conducted in accordance with the United States Public Health Service’s Policy on Humane Care and Use of Laboratory Animals. A 5 mm tail tip was sent to TransnetYX^®^, Inc (Cordova, TN) to confirm genotypes. Mice were group-housed according to sex and genotype on a 12:12 hour light / dark cycle, and laboratory rodent diet (LabDiet, 5001) and water were available *ad libitum*. One week after behavioral testing, mice were deeply anesthetized, weighed, and euthanized by decapitation. The brain was extracted and separated along midline. One hemibrain was fixed for immunofluorescent staining while the hippocampus and cortex from the other hemisphere were dissected on wet ice and stored at −80°C until processing. Subcutaneous, pgWAT, brown adipose tissue, and the liver were extracted by blunt tip dissection, individually weighed, and stored at −80°C until RT-qPCR analysis.

### Chemicals

Unless otherwise noted, all chemicals were prepared and stored according to manufacturer’s recommendations.

### Treatment

D-Met (Millipore Sigma; Cat: M9375) was dissolved in ddH_2_O and injected into sterilized water pouches every week for a final concentration of 2.67 mg/mL. This provided a daily dose of 400 mg/kg to mice by voluntary oral administration, as previously described ^9^. Due to variations in pathological burden and cognitive impairments across amyloidogenic models, treatment was initiated at 12-15 months in APP/PS1 mice and 18-20 months in APP^NL-F^ mice. Age- and sex-matched littermate controls received an equivalent volume of ddH_2_O into their water pouches as vehicle treatment. Cages were randomized to receive either D-Met or vehicle treatment. Table 1 provides the average ages of mice in each cohort when treatment started.

**Table 1:** Average age of each mouse cohort when treatment began.

| Cohort | Age (months) |
| --- | --- |
| Male APP/PS1 Vehicle | 13.39 ± 0.36 |
| Male APP/PS1 D-Met | 13.34 ± 0.26 |
| Male APP<sup>NL-F</sup> Vehicle | 19.89 ± 0.26 |
| Male APP<sup>NL-F</sup> D-Met | 19.27 ± 0.25 |
| Female APP/PS1 Vehicle | 13.43 ± 0.46 |
| Female APP/PS1 D-Met | 13.56 ± 0.52 |
| Female APP<sup>NL-F</sup> Vehicle | 19.54 ± 0.21 |
| Female APP<sup>NL-F</sup> D-Met | 19.76 ± 0.32 |

### Mouse water consumption and body weight

The amount of water consumed was determined weekly and normalized to the body weight of the mice in that cage. Water pouches with noticeable leaks were omitted from data analysis. Mice were weighed one week prior to treatment then weekly thereafter.

### Open field

After four weeks of D-Met treatment, ambulation and anxiety behavior were assessed in the open field test. The 16 x 16 x 16” ANY-Box (Stoelting Co.) open field maze consists of an unescapable opaque square wall-enclosure. After four weeks of D-Met treatment, mice enter the maze at the same corner and are given a 30 min exploration period. The ANY-maze video tracking system (Stoelting Co., Wood Dale, IL; RRID:SCR_014289) records distance traveled and duration along the periphery and center.

### MWM training and probe challenge

After five weeks of D-Met treatment, mice underwent cognitive assessment using the MWM spatial learning and memory recall paradigm, during which mice are trained to utilize visual cues to repeatedly swim to a static, submerged hidden platform. The MWM paradigm consisted of 5 consecutive learning days with three, 90-sec trials/day and a minimum 20-minute inter-trial interval. During the delayed memory recall, the platform was removed and mice were given a single, 60 second probe challenge. The ANY-maze video tracking system was used to record navigational parameters and data analysis. The three sessions for each training day were averaged for each mouse for analysis. Variables extracted from ANY-maze and utilized for data analysis include platform entries and latency, annulus 40 entries and latency, and swimming speed.

### mRNA extraction and quantification

mRNA expression was analyzed by quantitative RT–qPCR as previously described ^14^ using Bio-Rad cDNA synthesis kits (Cat # 1708897 and 1708891) and SYBR green mix (Cat # 1725121) and the QuantStudio 3 Real-Time PCR (Thermo Fisher Scientific). RNA was extracted using a RNeasy mini kit or RNeasy Lipid Tissue Mini Kit (Qiagen, Cat # 217004 and 74804) following the manufacturer’s instructions. Relative expression was calculated as previously described (Masternak *et al*., 2012) and primers were purchased from Integrated DNA Technologies. A list of forward and reverse primers used in this study is available in Table 2. Beta-2 microglobulin (B2M) and Ubiquitin conjugating enzyme E2D2 (UBE2D2) were used as housekeeping genes in pgWAT and hippocampus, respectively.

**Table 2:**
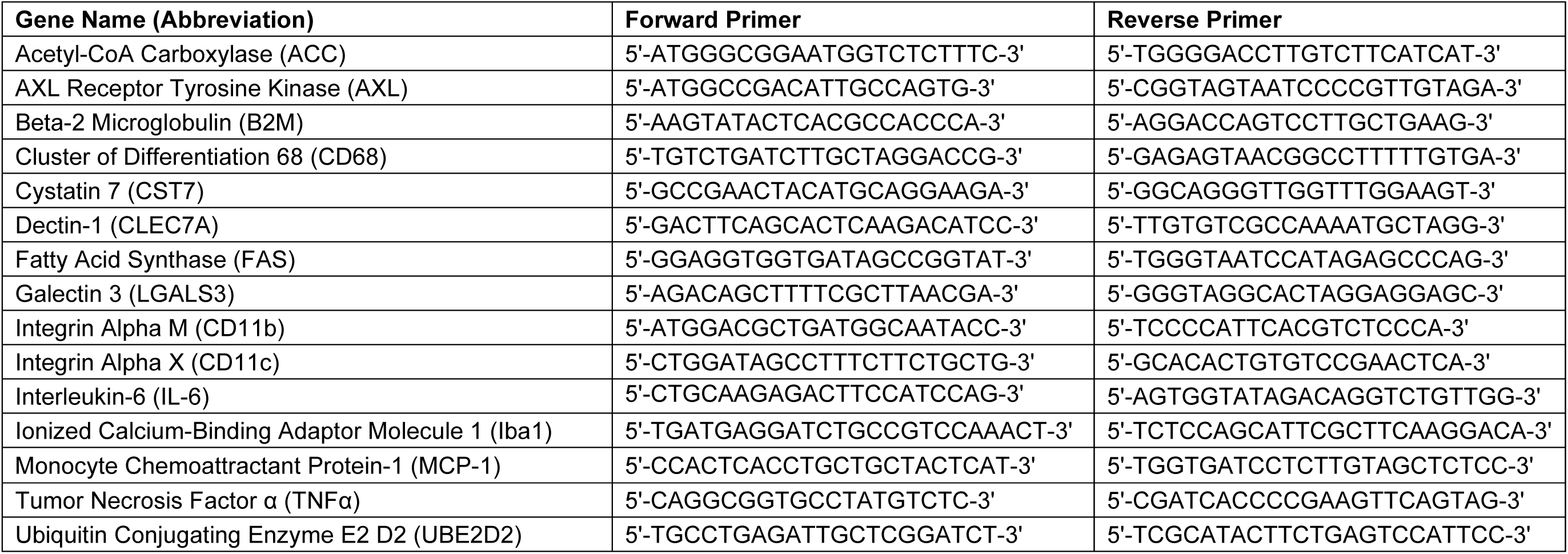
An alphabetized list of forward and reverse primers used in the present study.

### Amyloid plaque staining and semi-quantification

A hemibrain was fixed in 4% paraformaldehyde for 24-48 hrs then transferred into 30% sucrose in 0.1M phosphate buffer for at least 24 hours prior to sectioning. Twenty-micron coronal sections through the hippocampus were obtained using a CM1950 cryostat (Leica Biosystems). Every sixth serial section was stained for plaques using Amylo-Glo RTD with EtBr as a nucleic acid-specific fluorescing counter stain (1:10; Biosensis, Temecula, CA; Cat: TR-400-AG) according to the manufacturer recommended protocols. Stained slices were mounted and coverslipped using ProLong Gold Antifade Mountant (Thermo Fisher Scientific; Cat# P36934) ^15^. Imaging and quantification were conducted using a BZ-X810 fluorescent microscope (Keyence Corp.) with DAPI (Cat # OP-87762) and Texas Red (Cat # OP-87765) filters. The accompanying software was used to manually outline the hippocampus and retrosplenial cortex (RSC) and automatically determine the number of plaques in each brain region (mm^2^) and their size (µm^2^). Four hippocampal slices per mouse were averaged to obtain a single value for each subject. Amyloid plaques were identified by a dense spherical core of intense staining surrounded by a less compact spherical halo.

### Soluble Aβ_42_, lipid peroxidation, and glutathione reductase (GR) ELISA

The cortex from one hemisphere was dissected and stored at −80°C until tissue processing. Soluble Aβ_42_ concentrations were determined using the Human / Rat β amyloid(42) High Sensitive ELISA kit (WAKO Chemicals; Cat: 292-64501) according to the manufacturer recommended protocols. Cortical lipid peroxidation was determined by using the thiobarbituric acid reactive substances (TCA Method) assay (Cayman Chemicals; Cat: 700870) according to the manufacturer recommended protocols. Cortical glutathione reductase activity was determined by measuring the rate of NADPH oxidation in the sample using a GR assay kit (Cayman Chemicals; Cat: 703202).

### Statistical analysis

Prism software Version 10.6 (GraphPad Software, Inc., La Jolla, CA; RRID:SCR_002798) was used for all statistical analyses. A two-way ANOVA with a Fisher’s LSD post-hoc was used to test for significance of D-Met treatment within a sex and genotype for the body weight and MWM learning sessions. D-Met treatment differences within the same sex and genotype were determined using a one-tailed Student’s t-test for all remaining assays since our null hypothesis was predicting improvements compared to vehicle treatment. Potential outliers were determined with a single Grubb’s test (α=0.05). Individual mouse values are shown on all bar graphs and data are represented as mean ± SEM. Significance was defined as p<0.05.

## Results

### Weekly water consumption

All mice received either D-Met or vehicle in their drinking water. Daily consumption of D-Met was comparable to mice receiving vehicle (Figure 1A). No changes in drinking patterns were observed over the six weeks within treatment groups and across genotypes and sexes.

**Figure 1.**
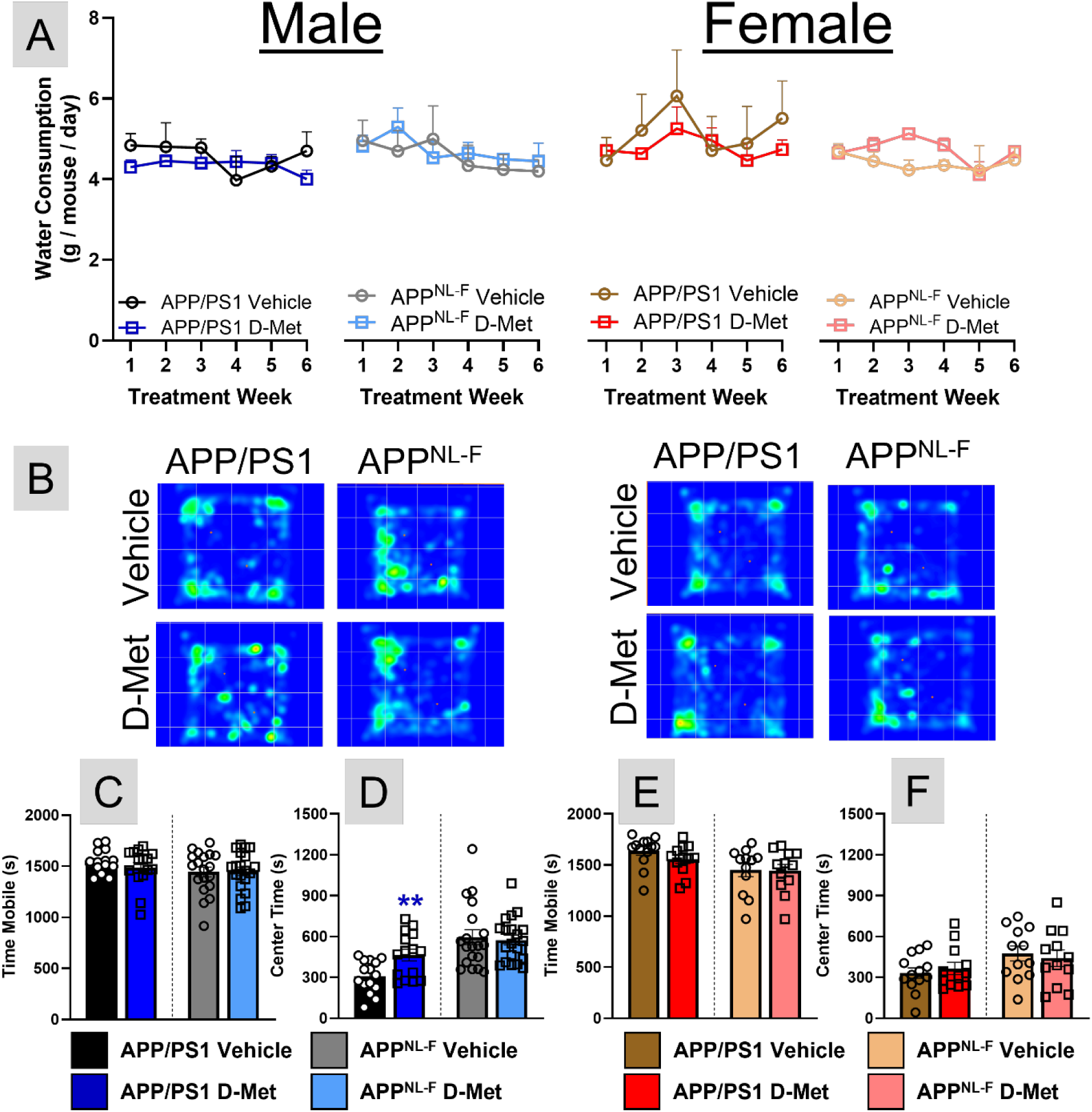
Water consumption (A) was calculated weekly for each mouse cage (n=3-5 cages per genotype and sex). Open field representative heat maps (B) are shown for each treatment, genotype, and sex. The time mice spent exploring the open field arena and its center are shown for male (C-D) and female (E-F) mice. A two-way ANOVA with Fisher’s LSD post hoc was used to compare weeks and treatment effects for all sexes and genotypes (A). A one-tailed Student’s t-test was used to assess D-Met effects within a sex and genotype, and individual mouse values are plotted on each bar graph (C-F). **p<0.01

### Assessment of mobility and anxiety

AD is associated with a decline in gait ^16^ and an increased prevalence of anxiety ^17^. The open field maze measures the general ambulatory and anxiety levels of rodents. As a survival mechanism, mice avoid brightly lit open areas and initially travel around the periphery before exploring the center ^18^. Increased center exploration corresponds with less anxiety. Figure 1B shows representative heat maps for the open field assay. No treatment differences were observed for either sex or genotype for the amount of time mice spent exploring the arena (Figures 1C, E). Male APP/PS1 mice receiving D-Met increased their center exploration time compared with sex-and genotype-matched controls (Figure 1D), which indicates reduced anxiety. Treatment effects were not observed for female APP/PS1 nor either sex of the APP^NL-F^ mice (Figures 1D, F). These data indicate that D-Met does not broadly alter locomotor function nor exploratory drive, reducing the likelihood that subsequent cognitive outcomes are confounded by changes in activity or anxiety.

### Effects of D-Met on spatial learning and memory

Loss of spatial cognition is a hallmark symptom of AD. During the five MWM training days, the distance traveled to locate the hidden submerged platform decreased for all cohorts of mice (Figures 2A-B). Initially, male APP^NL-F^ mice receiving D-Met mice had improved learning compared with sex- and genotype-matched vehicle controls, but this was equivalent by the fifth training session similar to the other cohorts of mice. During the delayed probe challenge, no differences in swimming speeds were observed for D-Met treated mice compared with sex- and genotype-matched vehicle controls (Figures 2C, E). The amount of time spent in the former location of the hidden escape platform was increased in female APP/PS1 mice treated with D-Met, but these effects were not observed in the other cohorts of mice on D-Met treatment (Figures 2D, F). Next, we examined an area slightly larger than the former location of the escape platform referred to as the annulus 40 ^19^. After D-Met treatment, female APP^NL-F^ mice had increased number of annulus 40 crossings, female APP/PS1 spent more time in the annulus 40, and male APP/PS1 mice had a better path efficiency into the annulus 40 compared with sex- and genotype-matched vehicle treated mice (Figure 2G-L). Altogether, these findings demonstrate that D-Met treatment preferentially enhances hippocampal-dependent memory recall rather than learning with effects more robust in female mice.

**Figure 2:**
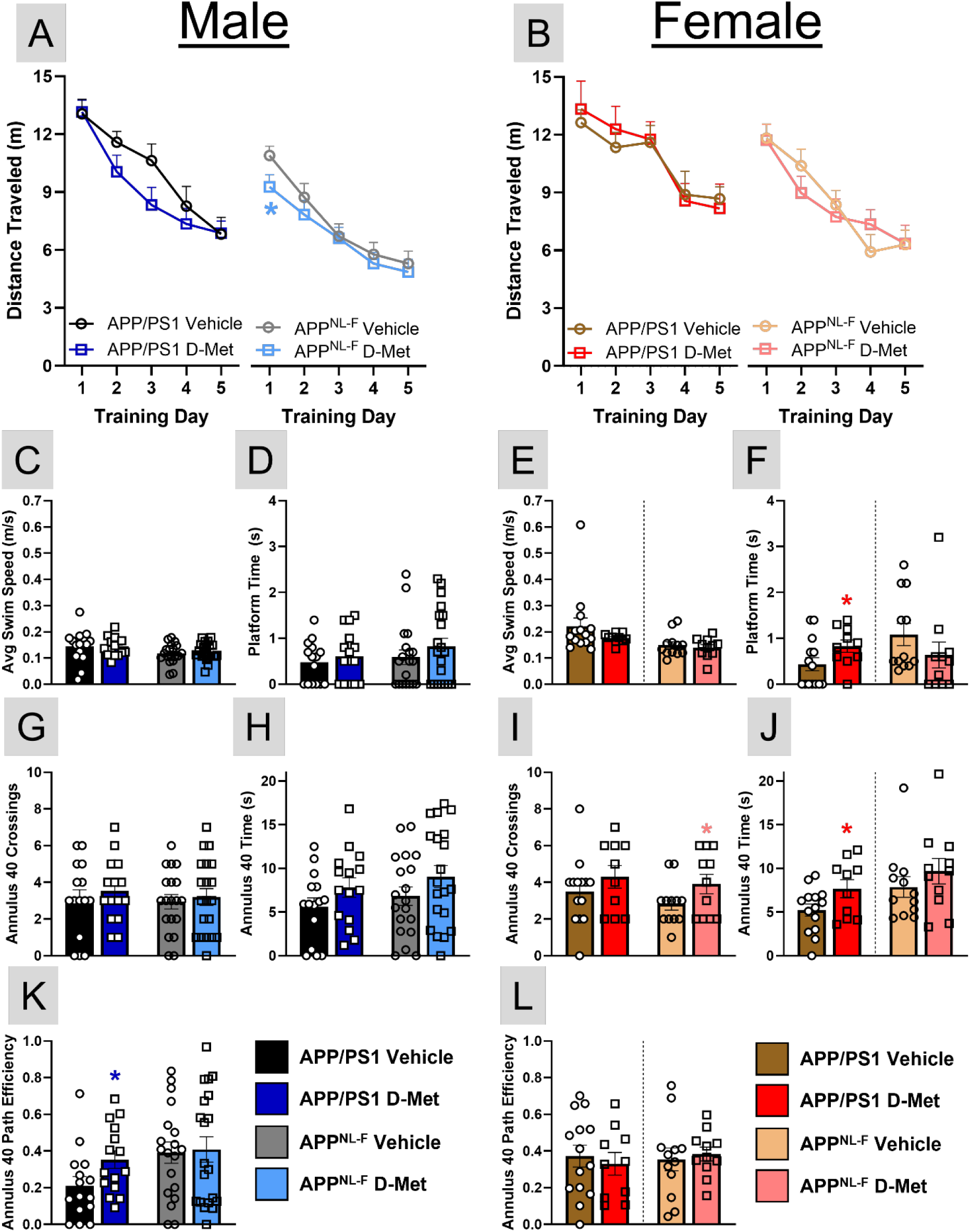
During the learning phase of the MWM, average distance traveled to reach the hidden escape platform during the five training days for male (A) and female (B) mice. Average speed, platform time, annulus 40 crossings, time, and path efficiency for male (C-D, G-H, K) and female (E-F, I-J, L) mice during the delayed probe challenge. A two-way ANOVA with Fisher’s LSD post hoc was used to compare training sessions and treatment effects (A-B) while a one-tailed Student’s t-test was used to measure D-Met effects within a sex and genotype (C-L). Individual mouse values are plotted on each bar graph. *p<0.05.

### Body weight changes during D-Met treatment and body composition post-treatment

Assessment of body composition revealed marked sex-specific effects of D-Met on body weight and adiposity. Male APP^NL-F^ exhibited increased body weight, subcutaneous adipose mass, and a pronounced increase in pgWAT (Figures 3A-B). Male APP/PS1 mice also demonstrated increased pgWAT mass, but to a lesser extent (Figure 3B). In contrast, female mice of either genotype showed no significant changes in body weight or fat depot mass (Figures 3A, C). No changes were observed in brown adipose tissue or liver weight in any group (Figures 3B-C). These results indicate that D-Met promotes adipose expansion selectively in males, particularly the APP^NL-F^ model.

**Figure 3:**
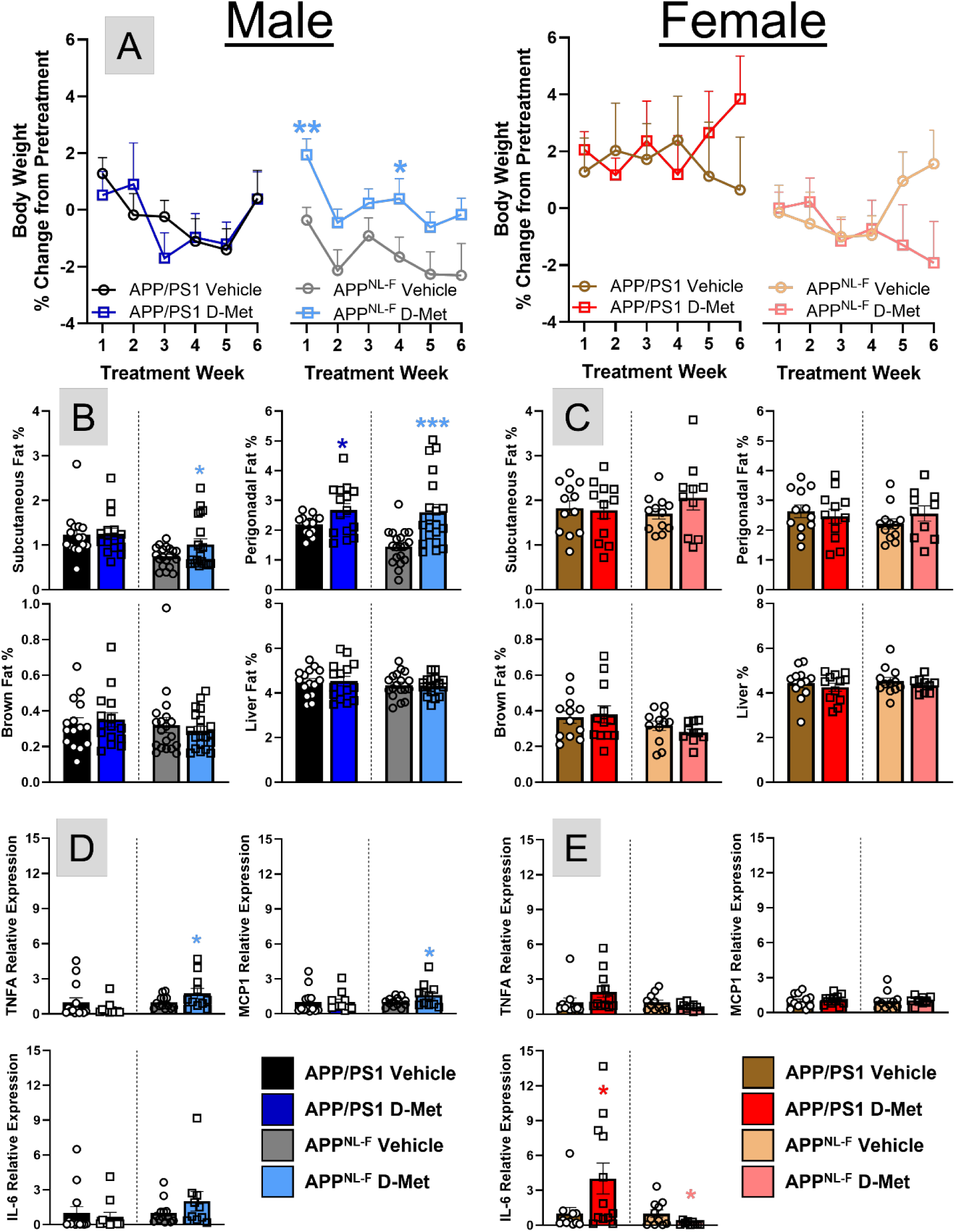
Mouse body weight was calculated weekly and the percent change from one week pretreatment was determined (A). The percent contribution of subcutaneous fat pgWAT, brown fat, and the liver to total body weight is shown for male (B) and female (C) mice. The relative expression of TNFα, MCP-1, and IL-6 against B2M in pgWAT are shown for male (D) and female (E) mice. A two-way ANOVA with Fisher’s LSD post hoc was used to compare weeks and treatment effects for all sexes and genotypes (A). A one-tailed Student’s t-test was used to assess D-Met effects within a sex and genotype, and individual mouse values are plotted on each bar graph (B-E). *p<0.05, **p<0.01, ***p<0.001.

### D-Met modulates adipose inflammatory signaling in a sex-dependent manner

To determine whether increased adiposity was associated with inflammatory remodeling, pgWAT was analyzed by RT-qPCR. Male APP^NL-F^ mice displayed elevated expression of TNFα and monocyte chemoattractant protein-1 (MCP-1), consistent with a pro-inflammatory adipose phenotype (Figure 3D). Surprisingly, these changes were not observed in male APP/PS1 mice after D-Met treatment. Female APP/PS1 mice exhibited increased IL-6 expression, whereas this expression was reduced in female APP^NL-F^ mice (Figure 3E). No changes were observed in genes regulating pgWAT mechanotransduction (piezo1 receptor; data not shown) or lipid synthesis pathways (acetyl-CoA carboxylase and fatty acid synthase; data not shown). These findings suggest that D-Met induces divergent adipose inflammatory responses, promoting inflammation in males while attenuating inflammatory signaling in female APP^NL-F^ mice.

### D-Met reduces amyloid pathology in a region- and sex-specific manner

Immunofluorescence was used to assay plaque burden in the hippocampus and RSC after D-Met treatment, two brain regions associated with spatial navigation and contextual memory. Representative whole hippocampal immunofluorescent images with 40x magnification insets to highlight plaque burden are shown in Figure 4A. Although no differences in plaque burden (number of plaques per mm of tissue) was observed in either brain region examined (data not shown), D-Met treatment reduced plaque size in the hippocampus of male APP^NL-F^ mice and the RSC of female APP/PS1 mice (Figures 4B-E). Soluble Aβ_42_ is the neurotoxic species associated with neuronal dysfunction and cognitive decline in AD ^20,21^. Soluble Aβ_42_ levels were decreased only in male APP/PS1 mice (Figures 5A-B). No global reduction in amyloid burden was observed across all groups after D-Met treatment. These data indicate that D-Met exerts modest but selective effects on amyloid pathology that depend on sex, genotype, and brain region.

**Figure 4:**
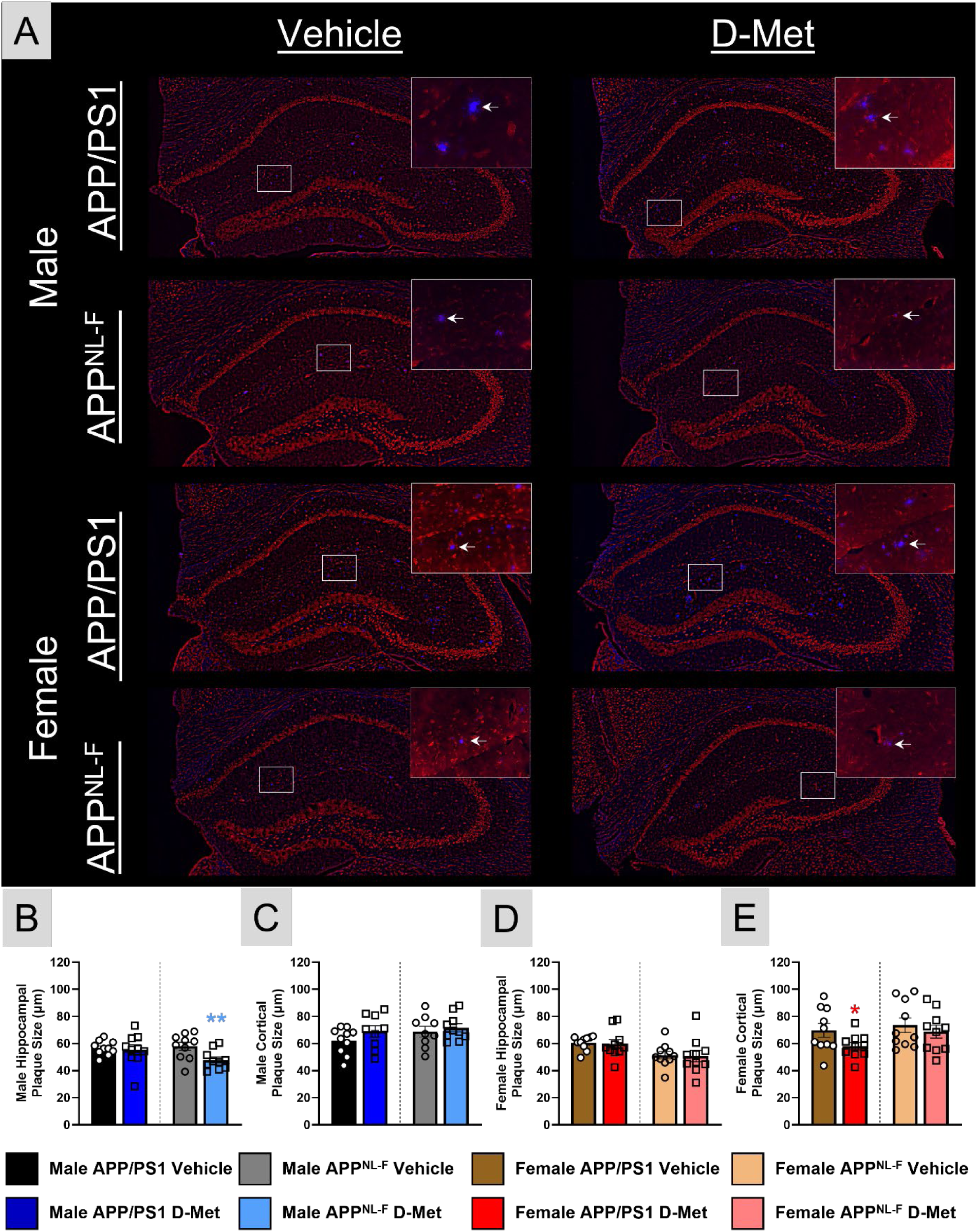
Representative immunofluorescent images (A) of whole hippocampal plaque burden (blue) counterstained with ethidium bromide (red). The inset is a 40x magnification of the white box in each whole hippocampal image. The (←) highlights an amyloid plaque. Average plaque size within the hippocampus and RSC for male (B-C) and female (D-E) mice. A one-tailed Student’s t-test was used to determine D-Met effects within each sex and genotype. Individual mouse values are plotted on each bar graph. *p<0.05, **p<0.01

**Figure 5:**
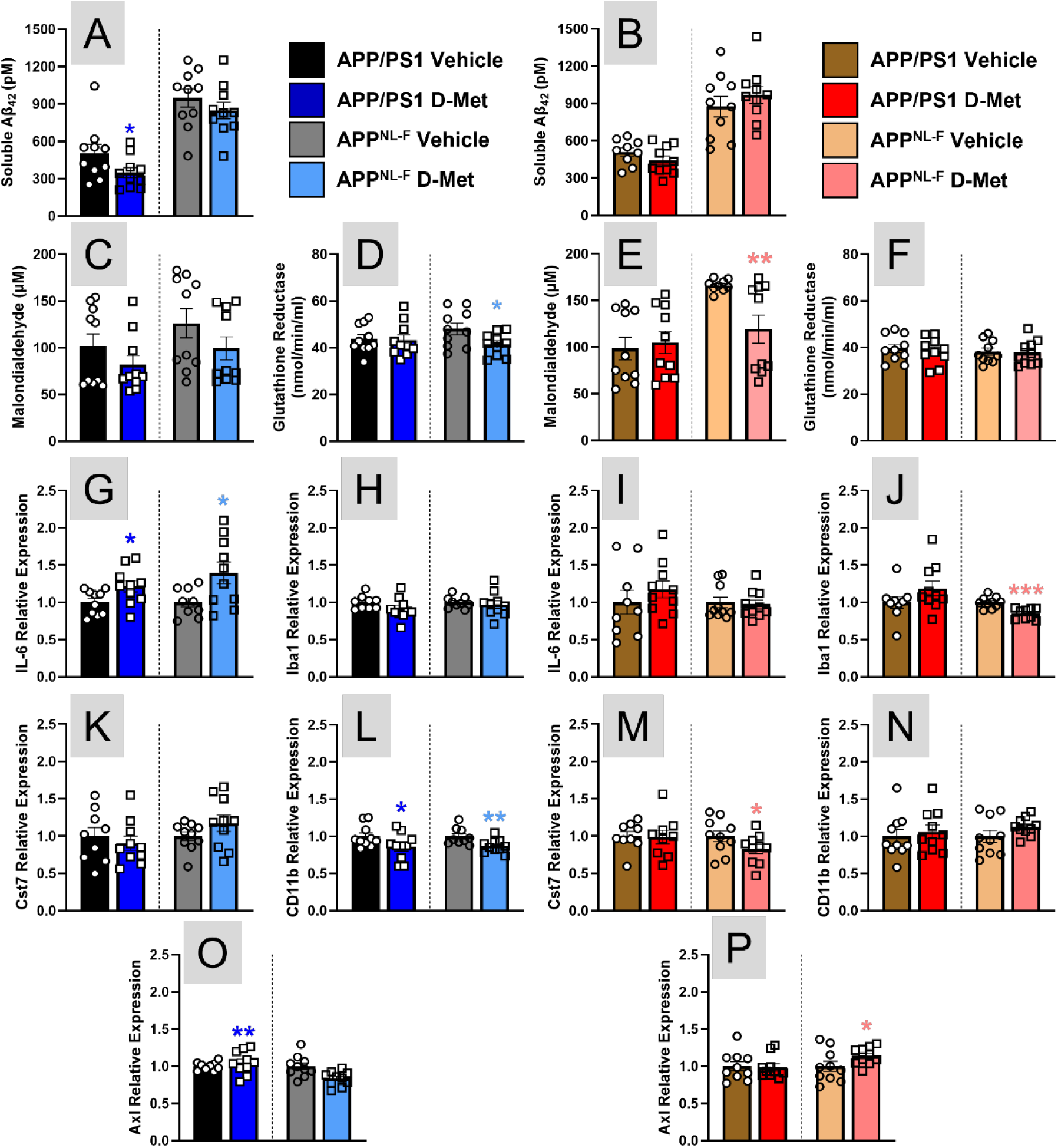
ELISA determination of soluble Aβ_42_ concentration, lipid peroxidation (MDA), and GR activity in the cortex of male (A, C, D) and female (B, E, F) APP/PS1 and APP^NL-F^ mice. Hippocampal relative expression of IL-6, Iba1, Cst7, CD11b, and Axl against UBE2D2 in male (G-H, K-L, O) and female (I-J, M-N, P) mice. A one-tailed Student’s t-test was used to determine D-Met effects within each sex and genotype. Individual mouse values are plotted on each bar graph. *p<0.05, **p<0.01, ***p<0.001

### D-Met attenuates oxidative stress primarily in female APP^NL-F^ mice

MDA is produced when lipids undergo peroxidation and is typically increased in AD. ELISA determination indicated that MDA was reduced in the cortex of female APP^NL-F^ mice receiving D-Met treatment compared with sex- and genotype-matched vehicle controls (Figures 5C, E). In contrast, GR levels were reduced in male APP^NL-F^ mice, with no changes observed in other groups (Figures 5D, F). These findings suggest that D-Met reduces lipid peroxidation most effectively in female APP^NL-F^ mice, aligning with the observed cognitive benefits in this cohort.

### D-Met differentially regulates neuroinflammatory and microglial signatures

Hippocampal inflammatory and microglial gene expressions were examined next. Hippocampal IL-6 expression following D-Met treatment was increased in both genotypes of male mice (Figures 5G), but TNFα expression was unchanged across all groups after D-Met treatment (data not shown). Notably, female APP^NL-F^ mice exhibited a marked reduction in Iba1 expression, accompanied by decreased expression of DAM marker Cst7 (Figures 5H, J, K, M). Cd11b expression was reduced in both male genotypes, while Axl expression was increased in male APP/PS1 and female APP^NL-F^ mice (Figures 5L, N, O, P). No changes were observed in CD68, Clec7a, Lgals3, or CD11c expression (data not shown). These data indicate that D-Met selectively remodels microglial phenotypes, with the most pronounced normalization occurring in female APP^NL-F^ mice that could further promote the spatial navigation improvements observed in this group.

## Discussion and conclusions

In the present study, we tested the hypothesis that short term oral D-Met administration would reduce amyloid burden and lipid peroxidation, thereby conferring disease-modifying benefits in mouse models of AD. The data partially support this hypothesis, revealing modest context-dependent cognitive benefits, primarily enhancing hippocampal spatial memory recall without altering exploratory or locomotor activity. The absence of changes in locomotor behavior support that D-Met improves hippocampal-dependent signaling.

These cognitive improvements are partially mediated by reductions in amyloid pathology and oxidative stress but are highly dependent on sex and genotype. Although D-Met reduced amyloid pathology in select regions, these changes were neither uniform nor sufficient to explain the observed behavioral improvements. This aligns with a growing consensus that amyloid load and cognitive performance can become uncoupled in later AD stages and cognitive outcomes may depend more on inflammatory mechanisms or individual resilience ^22^.

Rather, reductions in oxidative stress and microglial activation, most evident in APP^NL-F^ mice, may indicate a more relevant therapeutic mechanism. Oxidative stress is a well-established contributor to AD pathophysiology, with evidence that Aβ_42_ contributes to lipid peroxidation and neurodegenerative progression ^23^. This is typically mediated through ROS production and lipid peroxidation byproducts, such as MDA, that promote neuroinflammation – all of which are prevalent in animal models and post-mortem AD brains ^24,25^. This oxidative stress can precede plaque accumulation ^26^ and contribute to an excitotoxic environment that further drives neuronal loss ^21,27^.

Microglia play central roles in both amyloid clearance and generation of pro-oxidant species. Upon detection of rigid plaques, microglia become activated generating ROS and cytokines that contribute to a pro-inflammatory microenvironment ^28^. Our molecular data indicate that D-Met treatment remodels DAM gene expression, particularly in APP^NL-F^ mice. We observed reductions in Iba1 and Cst7 and enhanced Axl, a receptor tyrosine kinase that promotes phagocytosis while suppressing inflammation. The sex-specific nature of these effects is consistent with literature showing sexual dimorphic immunological responses and microglial metabolism in AD models, with female microglia showing distinct activation profiles compared to males ^29^.

Notably, D-Met induced substantial adipose expansion and inflammatory remodeling in male mice, particularly the APP^NL-F^ model. This peripheral inflammatory response may limit central benefits and highlights the influence of whole-body metabolic state on systemic inflammation, central immune responses, and cognitive outcomes. In contrast, female APP^NL-F^ mice, had reduced pgWAT IL-6 expression, lipid peroxidation, and microglial activation markers, which suggests a more favorable systemic-central interaction. Our laboratory has begun elucidating sex differences in metabolic and inflammatory phenotypes on cognition in response to glutamatergic modulation ^30^ and senolytic intervention ^19^. These sex differences in AD research are becoming increasingly common and underscore the need to consider sex as a biological variable in preclinical evaluation of therapeutic agents to improve personalized patient care.

Several study limitations should be acknowledged. Treatment duration was relatively short, and longer dosing paradigms may provide better disease-modifying effects. Additionally, direct measures of microglial phagocytic activity, inflammatory state, and Piezo1 signaling were not assessed, which limits mechanistic conclusions.

Collectively, these findings support a model where D-Met acts as a metabolic, redox, and microglial modulator rather than preventing plaque accumulation. Its therapeutic potential is most beneficial in the slower amyloid-progressing and lower inflammatory burden of the APP^NL-F^ model. This may suggest utilizing D-Met during the pre-symptomatic disease stage for optimal efficacy. Future studies incorporating microglial-specific functional assays may help to delineate the mechanisms underlying sex-specific responses while optimizing the therapeutic window for disease-modifying efficacy.

## Author contributions

MRP, JEC, TH, KQ, EDI, and AL conducted the experiments. MRP, EDI, and KNH analyzed the data. ERH, CB, and KNH conceived the study, designed the experiments, interpreted the data, and wrote the manuscript. All authors approved the final version of the manuscript.

## Statements and declarations

### Ethical considerations

Protocols for animal use were approved by the *Institutional Animal Care and Use Committee* at Southern Illinois University School of Medicine (Protocol number: 2025-055).

## Data availability

Data is available upon reasonable request.

## Abbreviations

Aβ: amyloid-β
AD: Alzheimer’s disease
APP: amyloid precursor protein
B2M: Beta-2 Microglobulin
D-Met: D-Methionine
DAM: disease-associated microglia
GR: glutathione reductase
pgWAT: perigonadal white adipose tissue
MCP-1: monocyte chemoattractant protein-1
MDA: malondialdehyde
MWM: Morris water maze
ROS: reactive oxygen species
RSC: retrosplenial cortex
UBE2D2: Ubiquitin Conjugating Enzyme E2 D2

